# Nonlinear EEG Analysis for Distinguishing Mind Wandering and Focused Attention: A Machine Learning Approach

**DOI:** 10.1101/2024.10.18.618974

**Authors:** Farshad Kheiri, Shyam Sundaram, Anatol Bragin

## Abstract

This study uses nonlinear analysis techniques to distinguish between mind wandering (MW) and focused attention (FA) states using EEG data. EEG recordings from 21 sessions were segmented into intervals of 2, 3, 5, 6, 10, and 15 seconds, and seven nonlinear features were extracted to capture the brain’s dynamic complexity. Machine learning models, including gradient boosting trees, were applied to classify MW and FA states, with the highest accuracy of 75% achieved using 5-second segments.

Frequency-related features, particularly mean frequency and global frequency, were the most important in distinguishing between MW and FA. These findings emphasize the role of nonlinear EEG analysis in understanding the chaotic brain patterns underlying cognitive states. Future work should focus on temporal dynamics and personalized models to improve classification accuracy, with potential applications in cognitive enhancement and mental health.

## Introduction

The brain is increasingly understood as a complex, nonlinear system capable of generating chaotic behavior, and nonlinear analysis techniques have gained significant attention in EEG data analysis. While linear EEG measures provide insights into the frequency domain correlating with mental states, they offer a limited perspective on the brain’s dynamic complexity and nondeterministic nature.

Nonlinear approaches, in contrast, offer a more advanced framework to explore the unpredictable and irregular patterns of neural activity underlying distinct cognitive states. Various nonlinear techniques have been employed to characterize EEG signals, offering a deeper insight into the chaotic dynamics associated with these states. For instance, entropy measures and fractal dimension analyses capture brain activity’s complexity. Entropy, specifically approximate entropy (ApEn) and sample entropy (SampEn), quantifies the level of randomness and disorder in EEG signals, directly measuring the system’s complexity.

Prior studies report reduced entropy during mind wandering (MW), signifying lower complexity compared to the more structured, less entropic state observed during focused attention (FA) (Elger et al., 2000; Natarajan et al., 2004; Paluš, 1996; Rodriguez-Bermudez & Garcia-Laencina, 2015). Fractal dimension (FD) analysis, which examines the self-similarity and scaling properties of EEG signals, also shows that MW is associated with more irregular, fractal-like signals, while FA induces a more homogenous, less complex pattern (Lu & Rodriguez-Larios, 2022; Natarajan et al., 2004; Zangeneh Soroush et al., 2018).

Higher-order spectral analysis, such as bispectral measures, extends Fourier analysis by considering the interactions between different frequency components in EEG signals. This approach is valuable for detecting the non-Gaussian and nonlinear characteristics of brain activity, which are crucial for differentiating MW from FA (Pradhan et al., 2012). Additionally, recurrence quantification analysis (RQA) and detrended fluctuation analysis (DFA) have demonstrated their potential in identifying long-range correlations and recurrence patterns in EEG signals that may distinguish MW from FA (Paluš, 1996; Rho et al., 2021; Wackermann, 1999).

Dimensional complexity estimates, such as those provided by the Grassberger-Procaccia algorithm, assess the geometric complexity of EEG trajectories in reconstructed state spaces, capturing intricate neural activity patterns (Elger et al., 2000; Lee et al., 2001; Natarajan et al., 2004; Paluš, 1996). Metrics like correlation dimension (CD) and Lyapunov exponents (LLE) quantify chaos and predictability in EEG data, revealing that MW is generally associated with a reduction in complexity compared to FA (Lu & Rodriguez-Larios, 2022). This indicates that FA entails more structured and ordered brain dynamics, while MW is characterized by more chaotic, unpredictable neural patterns, reflecting the spontaneous and internally driven nature of wandering thoughts (Lu & Rodriguez-Larios, 2022; Rho et al., 2021).

In this study, we apply multiple nonlinear techniques concurrently to compare the two mental states—mind wandering (MW) and focused attention (FA)—and evaluate their classification accuracy. Based on the existing literature, we have selected several EEG signal analysis methods representing power, frequency, and complexity across different segment durations and employed them in a nonlinear model to determine their efficacy in distinguishing MW from FA conditions.

## Methods

### Data acquisition

This study is part of a long-term single-participant EEG collection process. Data were acquired in a quiet room between 9 and 12 AM, with the participant seated comfortably in a chair. Recording sessions were spaced 2 to 3 days apart. EEG signals were collected using the Smarting Pro system (mBT amplifier, Serbia) with a sampling rate of 250Hz. A 32-channel 10×20 montage was applied using the Gelfree-S3 Cap (GreenTek). Prior to each experiment, electrode impedance was verified and maintained below 20 Ohms.

### Description of data acquisition from a first-person perspective

Each session consisted of 6 minutes of EEG recording with the participant’s eyes closed. A wood block click occurred three minutes into the session. The session began with 5 minutes of mind wandering (MW), followed by a shift to focused attention (FA) after the 50 dB wooden block click. During the MW phase, the participant mentally revisited past experiences, beginning with an imagined entry into their childhood home in western Siberia (Russia), allowing the mind to naturally reflect on various moments of life. Upon hearing the click, the participant switched to a focused breathing exercise, counting breaths from one to ten in a mindful breathing mode until the session concluded.

Following each session, the participant provided a brief written report describing their mind-wandering content. Additionally, the quality of the FA period was rated on a scale from five to one, where a score of five indicated no intrusive thoughts during the 3-minute FA session, 4.5 indicated one thought, and so on. No sessions recorded more than four thoughts during the FA phase. A total of 21 sessions were recorded in this series.

### Data analysis

Data was cleaned using EEGLAB 2023.0 (Delorme & Makeig, 2004). The following steps were executed: (1) importing raw data, (2) manually inspecting and removing electrostatic, muscle, and other behavioral artifacts, (3) applying spherical interpolation for any bad channels (typically one or two per recording) following EEGLAB protocols, (4) performing ICA decomposition, and (5) eliminating independent components (ICs) related to eye movement, muscle activity, and other artifacts. The transition between MW and FA was isolated using a 30-second window before and after the sound click in EEGLAB_2023 software.

### Non-linear approaches

A total of 42 recordings were analyzed, with 21 from the MW state and 21 from the FA state, each lasting 30 seconds. We applied the following non-linear measures to assess their sensitivity and specificity in distinguishing MW from FA mental states:

1. Global Field Power (GFP): GFP measures the overall strength of brain activity across all channels, reflecting the intensity of electrical signals. Higher GFP values indicate more intense brain activity.
2. Global Frequency (GFreq) GFreq identifies the dominant frequency of electrical activity across all channels, highlighting the prevailing brainwave types (e.g., theta, delta, beta, gamma) corresponding to different brain states.
3. Global Complexity (GComplexity): GComplexity assesses the unpredictability or intricacy of brain activity. High complexity values suggest diverse and dynamic brain states.
4. Mean Power Across Channels: This feature calculates the average signal strength across all channels, providing a sense of the typical energy level of the EEG signals.
5. Standard Deviation of Power Across Channels: This captures the variation in brain activity strength across different channels, with higher values indicating greater regional variability in brain activity.
6. Mean Frequency Across Channels: This metric gives the average rate of brainwave oscillations across all channels, offering insight into the overall pace of neural activity.
7. Standard Deviation of Frequency Across Channels: This feature measures how much brainwave oscillation rates vary across different channels, with greater variation suggesting differential brain region functioning.

Further details on these features are provided in Appendix A.

We applied several machine learning algorithms to analyze the data, including random forest, multi-layer perceptron (a neural network algorithm), support vector machines, gradient boosting trees, and logistic regression. The data were divided into training and test sets, with the test set comprising six randomly selected recordings (three from each mental state). (Figure 1.)

**Figure 1:**
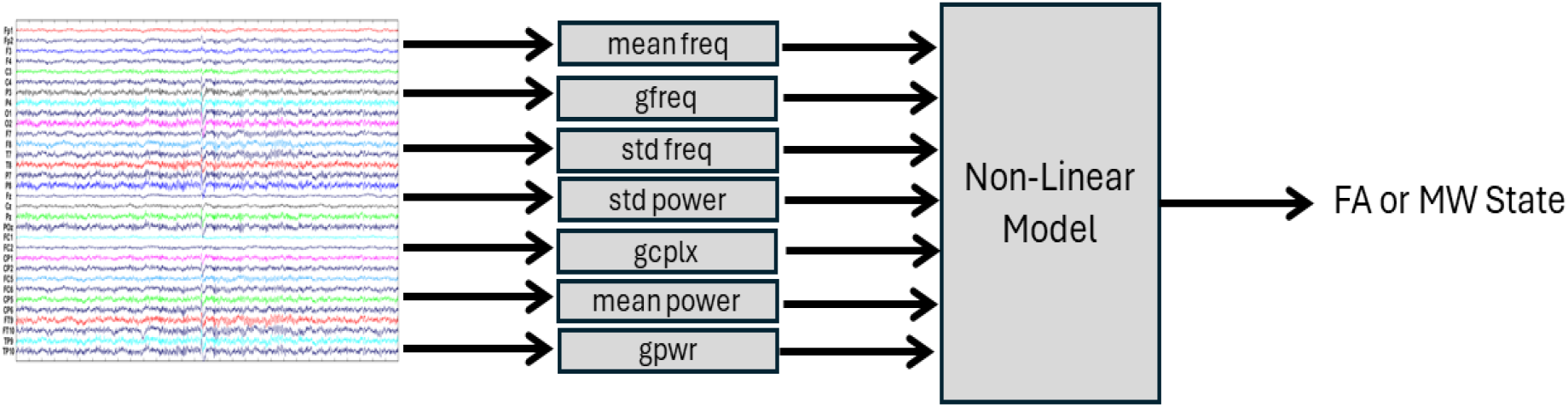
The classifier to distinguish between the two states (FA and MW).

## Results

Each EEG recording was segmented into multiple time intervals, and the seven nonlinear features described in the feature engineering section were applied to compare the states of mind wandering (MW) and focused attention (FA). The segment lengths were 2, 3, 5, 6, 10, and 15 seconds. Subsequently, machine learning models were trained using each of the seven nonlinear features, and 10-fold cross-validation was employed to mitigate the risk of overfitting.

Among the various models tested, the gradient boosting trees algorithm demonstrated superior performance compared to other techniques. The highest accuracy on the test set was achieved using 5-second segments, with an accuracy of 75%. This means that, initially, each recording was divided into 5-second intervals, and the model was applied to these shorter segments. Therefore, instead of working with a single 30-second MW or FA segment, each recording was treated as six distinct 5-second MW or FA states (see Table 1).

**Table 1.** Accuracy of the gradient boosting trees model for predicting MW and FA states across different segment durations.

| Segment Length (sec) | Accuracy on the Testing Set |
| --- | --- |
| 2 | 55.56% |
| 3 | 51.67% |
| 5 | 75% |
| 6 | 53.33% |
| 10 | 61.11% |
| 15 | 63.90% |

In addition to evaluating the overall accuracy, the importance of each engineered feature was analyzed to determine which feature contributed most to distinguishing between MW and FA states in the 5-second segments. The analysis revealed that the mean frequency (mean_freq) was the most influential feature, followed by global frequency (gfreq) and standard deviation of frequency (std_freq), as detailed in Table 2.

**Table 2.**
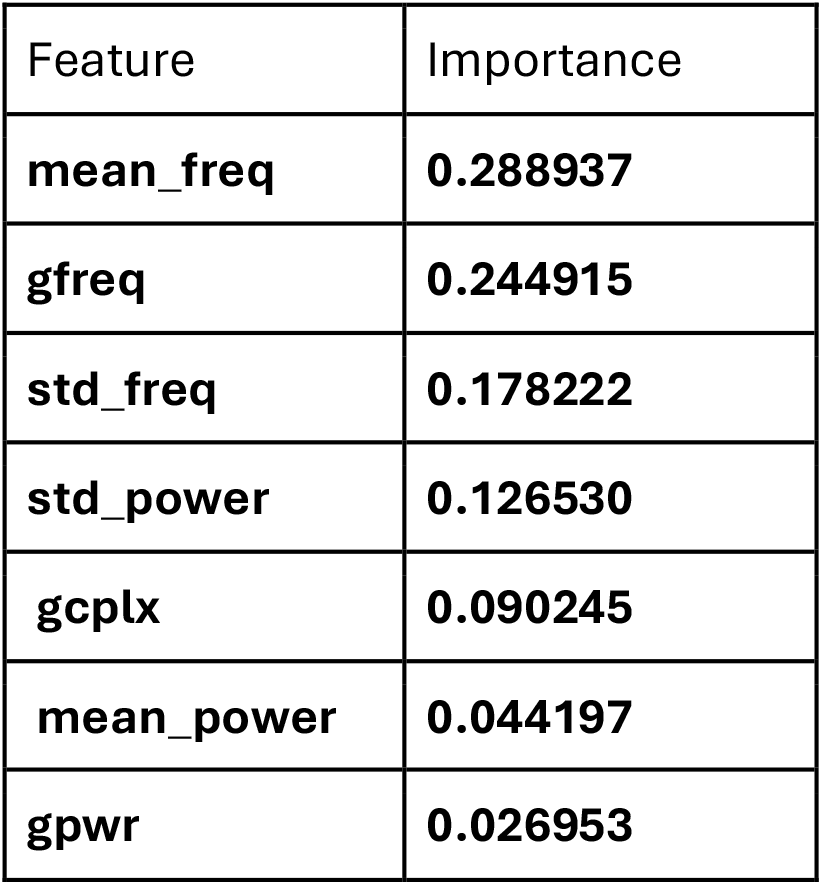
Feature importance in the gradient boosting trees model for 5-second segments.

## Conclusion

This study explored the application of nonlinear techniques in distinguishing between mind wandering (MW) and focused attention (FA) states using EEG recordings. By employing a diverse range of nonlinear features—representing power, frequency, and complexity— across varying segment durations, we aimed to assess the classification accuracy of these features in identifying different mental states. Among the models tested, gradient boosting trees outperformed other machine learning techniques, achieving the highest accuracy of 75% with 5-second segments. This segmentation approach allowed for a more granular analysis, treating each 5-second interval as an independent representation of MW or FA. This approach could be extended to the identification of other mental states.

The results demonstrated that frequency-based features, particularly mean frequency (mean_freq) and global frequency (gfreq), were the most influential in distinguishing between MW and FA. This suggests that the dynamic changes in brainwave frequencies are key markers of these mental states. Moreover, the analysis highlighted the importance of segment duration, with shorter segments (5 seconds) providing more accurate classifications than longer ones. This implies that temporal variations in cognitive states may be better captured in brief time windows.

Overall, these findings contribute to our understanding of the brain’s nonlinear dynamics during different cognitive processes. They underscore the potential of nonlinear EEG analysis to reveal the complex, chaotic patterns of neural activity underlying distinct mental states. Future research should investigate the underlying mechanisms that drive these frequency changes and explore the broader applicability of these nonlinear techniques across various cognitive tasks. Additionally, refining machine learning models to leverage these nonlinear features better could further enhance the accuracy of mental state classification in EEG studies.

### Next Steps

Future research should pursue several directions to advance the classification of mind wandering (MW) and focused attention (FA) states. First, the role of temporal dynamics warrants further investigation. The effectiveness of shorter time segments suggests that time-dependent patterns in brain activity could be better captured by models that account for these dynamics, such as recurrent neural networks (RNNs) or long short-term memory (LSTM) networks. These deep learning algorithms are well-suited for handling sequential data and may enhance the model’s ability to track cognitive state transitions over time.

Second, the variability observed in feature importance across different channels points to the potential value of individualized models. Neural correlates of MW and FA may differ across individuals, and personalized models that account for unique brain activity patterns could improve classification accuracy. Such an approach would better tailor the analysis to each individual’s neural signature.

Lastly, examining the transitions between MW and FA states using time-series models could provide deeper insights into how the brain navigates between introspective and focused modes of attention. Understanding these dynamic transitions in real time could contribute to more nuanced theories of cognitive state regulation and offer new possibilities for enhancing focus or managing wandering thoughts.

This research lays an essential foundation for future work to refine our understanding of brain states. By exploring the temporal dynamics, individual variability, and transitional patterns between cognitive states, we can move closer to developing targeted applications for cognitive enhancement and mental health interventions.

## Acknowledgements

For help with this writing, we used the following AI tools: https://notebooklm.google/

## Appendix A.

Detailed features of non-linear parameters

1. Global Field Power (GFP):
  - Mathematical Formula:

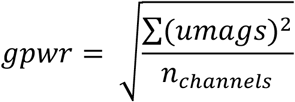

where *umags* represents the magnitudes of the EEG signal for each channel, and *n*_*channels*_ is the total number of EEG channels.
2. Global Frequency (GFreq):
  - Mathematical Formula:

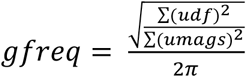

where *udf* represents the first differences (the rate of change between consecutive time points) of the EEG signal and *umags* are the magnitudes of the EEG signal.
3. Global Complexity (GComplexity):
  - Mathematical Formula:

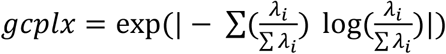

where *λ*_*i*_ are the eigenvalues of the covariance matrix of the EEG signals from all channels.
4. Mean Power Across Channels:
  - Mathematical Formula:

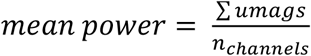

where *umags* are the magnitudes of the EEG signals, and *n*_*channels*_ is the total number of EEG channels.
5. Standard Deviation of Power Across Channels:
  - Mathematical Formula:

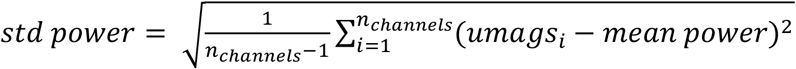

where *umags* are the magnitudes of the EEG signal, and *n*_*channels*_ is the total number of channels.
6. Mean Frequency Across Channels:
  - Mathematical Formula:

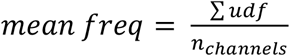

where *udf* represents the rate of change of the EEG signal, and *n*_*channels*_ is the total number of channels.
7. Standard Deviation of Frequency Across Channels:
  - Mathematical Formula:

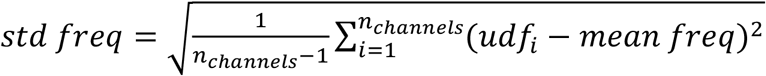

where *udf* represents the rate of change of the EEG signal, and *n*_*channels*_ is the total number of channels.

